# Multi-omics characterization of vascular, neurodegenerative, and mixed neuropathology in the aging human brain

**DOI:** 10.64898/2026.03.17.712511

**Authors:** Annie J. Lee, Minghua Liu, Elanur Yilmaz, Caghan Kizil, Shahram Oveisgharan, Julie A. Schneider, David A. Bennett, Richard Mayeux, Badri N. Vardarajan

**Affiliations:** Department of Neurology, College of Physicians and Surgeons, Columbia University Irving Medical Center, New York, NY 10032, USA; Taub Institute for Research on Alzheimer’s Disease and the Aging Brain, College of Physicians and Surgeons, Columbia University Irving Medical Center, New York, NY 10032, USA; The Gertrude H. Sergievsky Center, College of Physicians and Surgeons, Columbia University Irving Medical Center, New York, NY 10032, USA; Rush Alzheimer Disease Center, Rush University Medical Center, Chicago, IL 60612, USA

## Abstract

Late-life cognitive impairment most commonly occurs in the setting of mixed neurodegenerative and cerebrovascular pathology, yet the molecular programs distinguishing vascular, neurodegenerative, and mixed pathology in the aging human brain remain incompletely defined. We performed neuropathology-stratified proteomic and transcriptomic profiling of postmortem brain tissue from participants in the Religious Orders Study and Rush Memory and Aging Project. Dorsolateral prefrontal cortex proteomics (n = 733) were analyzed alongside bulk RNA sequencing from dorsolateral prefrontal cortex (n = 938), posterior cingulate cortex (n = 569), and anterior caudate (n = 632). Participants were classified into vascular, neurodegenerative, and mixed pathology groups based on comprehensive autopsy assessment. Neurodegenerative and mixed pathology, relative to vascular pathology, showed coordinated upregulation of immune and inflammatory pathways and downregulation of mitochondrial and oxidative phosphorylation programs across molecular layers. Although few individual proteins differed between mixed and neurodegenerative groups, pathway-level analyses identified additional remodeling programs in mixed pathology, including extracellular matrix organization and vesicle-mediated transport. Proteomic co-expression network analysis identified immune–stress modules associated with amyloid burden, tau pathology, and cognitive decline, whereas a mitochondrial bioenergetic module showed relative preservation in vascular pathology. Cross-omics concordance was robust at the pathway level but limited at the level of individual genes and proteins. These findings define conserved molecular programs distinguishing vascular, neurodegenerative, and mixed pathology and demonstrate that pathway-level organization provides a stable framework for interpreting molecular heterogeneity in late-life dementia.

## INTRODUCTION

Late-life cognitive impairment arises within a neuropathologic landscape characterized by substantial biological heterogeneity. Although Alzheimer’s disease (AD) is the most common neurodegenerative disorder worldwide, autopsy studies consistently demonstrate that age-related brain pathologies rarely occur in isolation. Neurodegenerative proteinopathies—including amyloid-β plaques, tau tangles, α-synuclein inclusions, and TDP-43 pathology—frequently coexist with cerebrovascular lesions such as macroinfarcts, microinfarcts, arteriolosclerosis, and cerebral amyloid angiopathy(*1*). In community-based cohorts, 30–50% of individuals meeting clinicopathologic criteria for AD exhibit substantial cerebrovascular pathology (*1–4*).

The Religious Orders Study and the Rush Memory and Aging Project (ROSMAP) have provided detailed longitudinal clinicopathologic characterization of this complexity. In this cohort, more than 80% of older individuals harbor mixed degenerative and vascular lesions at autopsy(*5*). One analysis identified over 230 unique combinations of coexisting pathologies across participants, underscoring the combinatorial diversity of age-related brain disease. Importantly, these mixed pathologies cluster into structured neuropathologic profiles associated with markedly different cognitive trajectories(*5*). Mixed pathology therefore represents the prevailing biological state of the aging brain.

Within ROS/MAP, neuropathologic stratification has defined vascular, neurodegenerative, and mixed pathology groups based on the presence or absence of substantial cerebrovascular and neurodegenerative lesions(*6*). In this cohort, cerebrovascular disease pathologies—including macroinfarcts, microinfarcts, and arteriolosclerosis—are independently associated with cognitive impairment(*6*). Macroinfarcts, particularly in frontal white matter, are linked to faster cognitive decline in individuals without substantial Alzheimer disease neuropathologic change (ADNC) or other major neurodegenerative pathologies. In contrast, high burdens of ADNC and limbic-predominant age-related TDP-43 encephalopathy neuropathologic change, often accompanied by hippocampal sclerosis, are associated with the steepest longitudinal cognitive trajectories (*5*). These observations establish that vascular and neurodegenerative pathologies contribute differentially to clinical expression and frequently coexist within the same brain. Cerebrovascular pathology is also associated with structural brain changes, including white matter injury and increased atrophy(*7*). Pathologic studies link large and small vessel disease to impaired cerebral blood flow and blood–brain barrier abnormalities, features commonly observed in aging brains with mixed pathology(*7*).

The genetic architecture of late-onset AD intersects with vascular biology. The *APOEε4 a*llele—the strongest risk locus for late-onset AD—associates not only with amyloid deposition but also with cerebral amyloid angiopathy and small vessel disease(*8*). Additional risk loci, including *ABCA7, CLU, CR1, PICALM,* and *SORL1*, implicate lipid metabolism, innate immune signaling, and endosomal trafficking pathways with relevance to both neurodegenerative and cerebrovascular processes(*9–13*). Clinicopathologic stratification and genetic studies indicate that vascular and neurodegenerative mechanisms converge at shared biological pathways. However, whether neuropathologically defined vascular, neurodegenerative, and mixed groups are reflected in distinct molecular programs within the aging human brain remains unknown.

Here, leveraging deeply phenotyped participants from ROSMAP, we performed a neuropathology-stratified cross-modal analysis of postmortem human brain tissue. We analyzed dorsolateral prefrontal cortex proteomics alongside bulk RNA sequencing across multiple cortical and subcortical regions and evaluated concordance at the gene and pathway levels across molecular layers. This design enabled us to define coordinated biological programs associated with neurodegenerative, vascular, and mixed pathology and to assess their conservation across brain regions.

## RESULTS

### Cohort and data description

We performed an in-depth multi-omic characterization of postmortem human brain tissue from participants in the Religious Orders Study and Memory and Aging Project (ROSMAP) to identify molecular programs underlying neurodegenerative, vascular, and mixed pathology. Participants were classified into three neuropathologically defined and mutually exclusive groups — neurodegenerative pathology, vascular pathology, and mixed pathology—based on comprehensive postmortem assessments of neurodegenerative and vascular pathological features (*6*) (**Table 1**). Proteomic profiling of the dorsolateral prefrontal cortex (DLPFC) was performed in 733 participants with 10,030 proteins quantified. Transcriptomic analyses were conducted across three brain regions, including the dorsolateral prefrontal cortex (DLPFC) in 938 participants with 18,629 transcripts, the posterior cingulate cortex (PCC) in 569 participants with 19,017 transcripts, and the anterior caudate (AC) in 632 participants with 17,574 transcripts. All participants in the neurodegenerative and mixed pathology groups had a pathological diagnosis of Alzheimer’s disease, whereas pathological AD was absent in the vascular pathology group. Subgroup distributions were consistent across omics layers. Participant demographic and clinical characteristics by omics modality and pathology subgroup are summarized in **Table 1**.

**Table 1.**
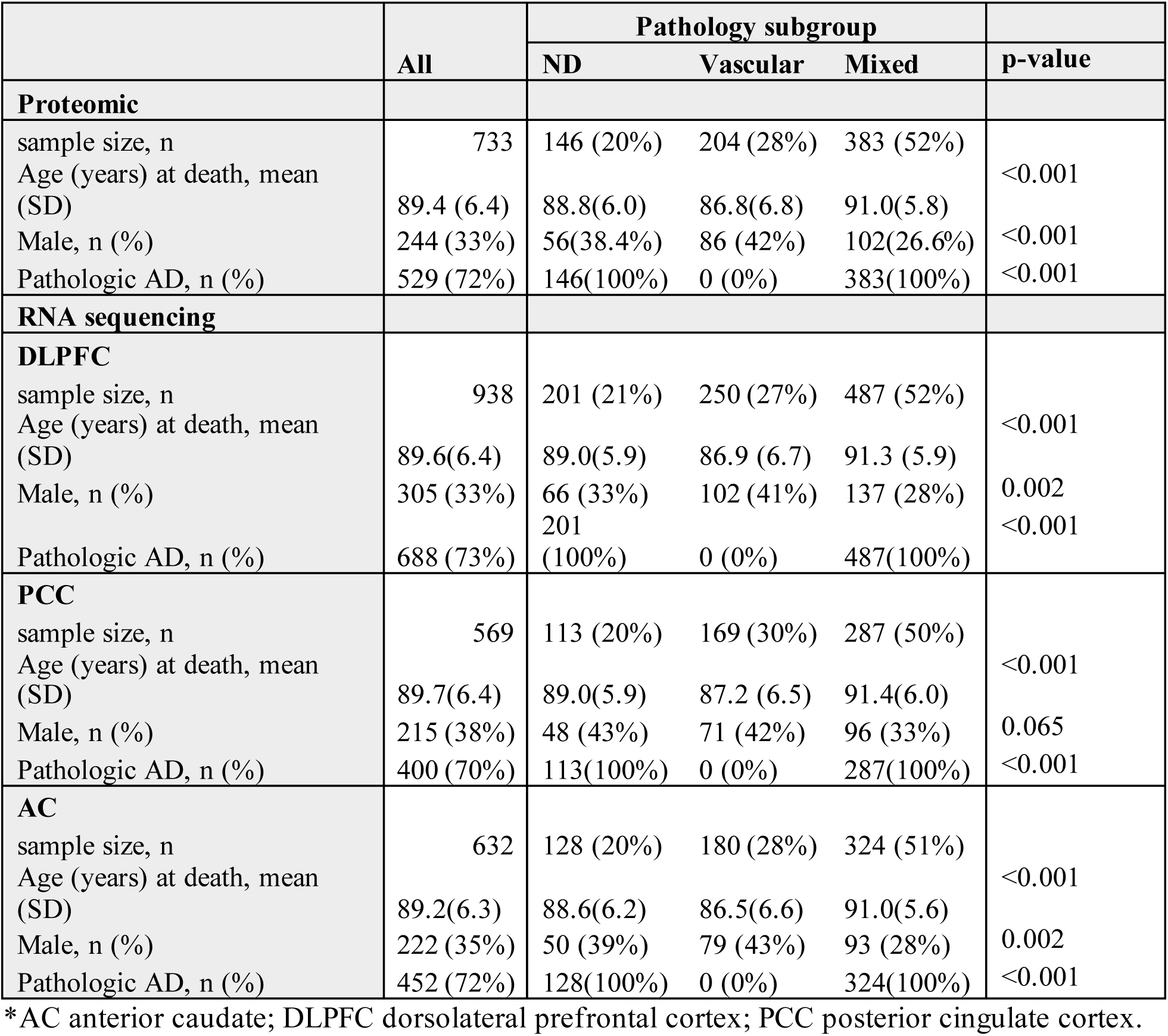
Characteristics of ROSMAP participants by neuropathology group.

### Proteomics analysis reveals shared immune activation and mitochondrial suppression in neurodegenerative and mixed pathology relative to vascular pathology

To identify molecular programs distinguishing neurodegenerative, vascular, and mixed pathology, we performed differential protein expression analysis in DLPFC for three pairwise contrasts—neurodegenerative versus vascular, mixed versus vascular, and mixed versus neurodegenerative—with adjustment for age, sex, and technical covariates. A large number of differentially expressed proteins (DEPs) were identified when comparing neurodegenerative pathology with vascular pathology (n = 1,006 DEPs) and when comparing mixed pathology with vascular pathology (n = 1,797 DEPs) (FDR < 0.05) (**Table S1** and **S2**). In contrast, no proteins were significantly differentially expressed between mixed and neurodegenerative pathology, indicating broad molecular similarity between these two groups (**Fig. 1A** and **1B**, **Table S3**).

**Fig. 1.**
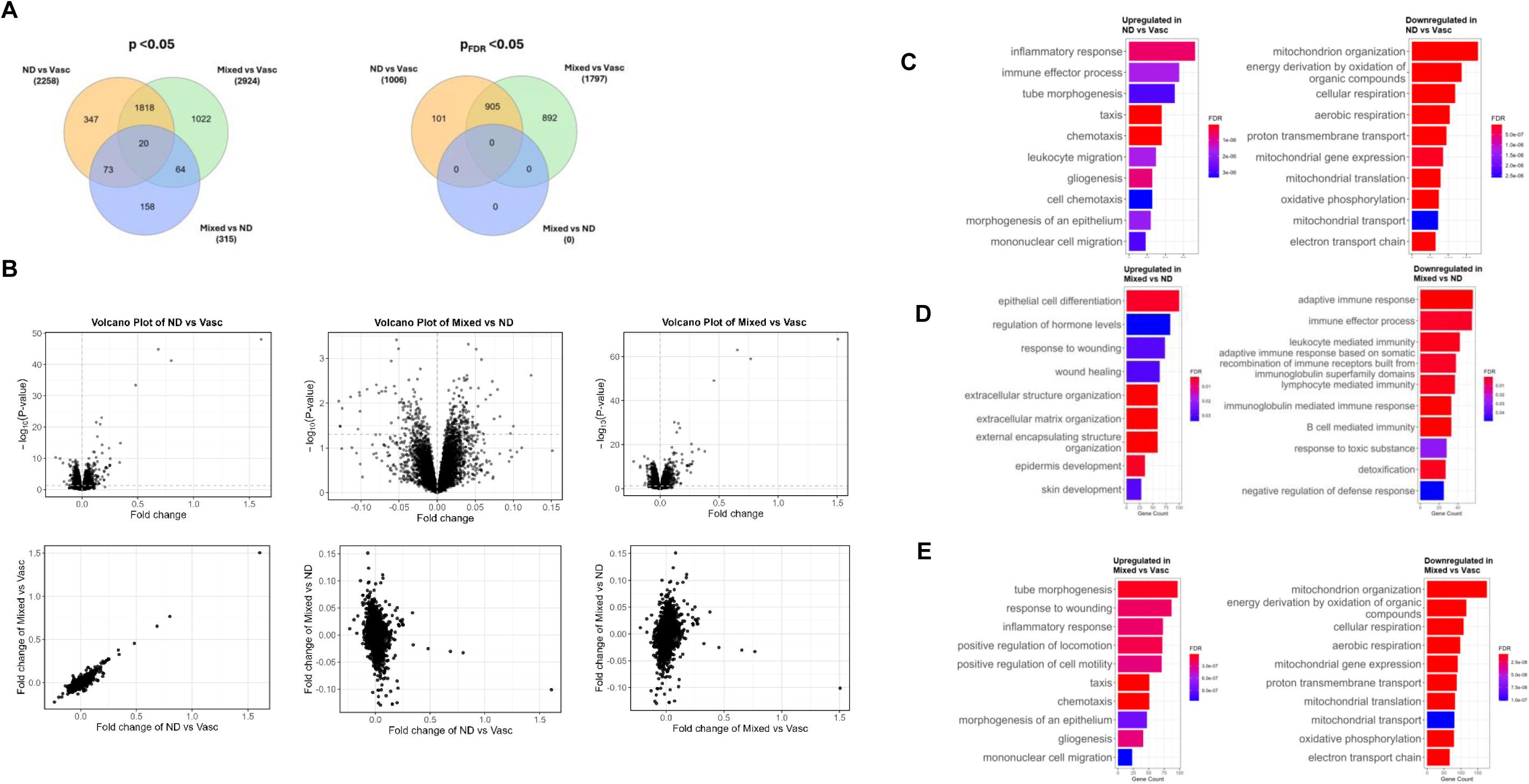
Proteomic differences across neuropathology groups. (**A**) Overlap of differentially expressed proteins across neurodegenerative versus vascular, mixed versus vascular, and mixed versus neurodegenerative comparisons (p < 0.05 and FDR < 0.05). (**B**) Volcano plots (top) and scatter plots of log fold-changes between comparisons (bottom). (**C–E**) Gene Ontology enrichment analyses for neurodegenerative versus vascular, mixed versus vascular, and mixed versus neurodegenerative comparisons. Differential expression was assessed using linear regression adjusted for age, sex, and technical covariates; FDR < 0.05.

Gene set enrichment analysis using Gene Ontology Biological Process annotations revealed extensive pathway-level differences in the neurodegenerative–vascular comparison, comprising 174 upregulated and 46 downregulated pathways (FDR < 0.05; **Table S4**). Upregulated pathways were predominantly enriched for immune and inflammatory processes, including leukocyte migration, chemotaxis, immune effector activity, angiogenesis, and gliogenesis. In contrast, downregulated pathways converged on mitochondrial organization, oxidative phosphorylation, cellular respiration, and broader bioenergetic functions (**Fig. 1C**).

A highly concordant enrichment profile was observed in the mixed versus vascular comparison. Upregulated pathways were similarly dominated by immune and inflammatory processes, whereas downregulated pathways were enriched for mitochondrial and bioenergetic functions (176 upregulated and 44 downregulated pathways; FDR < 0.05) (**Fig. 1D** and **Table S5**). The neurodegenerative–vascular and mixed–vascular contrasts showed strong concordance at the pathway level, sharing 158 upregulated pathways (Jaccard index = 0.82; *p* = 1.6 × 10⁻¹⁵) and 38 downregulated pathways (Jaccard index = 0.73; *p* = 1.0 × 10⁻⁴) (**Table S6**). In addition to this shared immune–metabolic signature, mixed pathology exhibited further enrichment of structural and tissue remodeling pathways, including response to wounding and tube and epithelial morphogenesis, consistent with enhanced vascular and cellular reorganization superimposed on a shared inflammatory program (**Fig. 1D**).

Pathway-level analysis revealed coordinated molecular differences between mixed and neurodegenerative pathology, despite minimal differences at the single-protein level (**Fig. 1E**). Mixed pathology showed enrichment of extracellular matrix organization, epithelial differentiation, wound healing, and tissue remodeling pathways, whereas neurodegenerative pathology was characterized by stronger enrichment of adaptive immune processes, including immunoglobulin-and lymphocyte-mediated immune responses (**Fig. 1E** and **Table S7**). These results indicate that, despite broad proteomic similarity, mixed pathology engages additional structured remodeling programs that distinguish it from neurodegenerative pathology alone.

### Proteomic co-expression networks identify shared immune–mitochondrial modules and mixed-specific remodeling

To characterize coordinated proteomic networks underlying neurodegenerative, vascular, and mixed pathology, we performed weighted gene co-expression network analysis (WGCNA) on DLPFC proteomic data. Of the 21 co-expression modules identified, six showed significant associations with clinicopathological traits, as well as with subgroup comparisons distinguishing neurodegenerative–vascular and mixed–vascular pathology (**Fig. 2A** and **Fig. S1**).

**Fig. 2.**
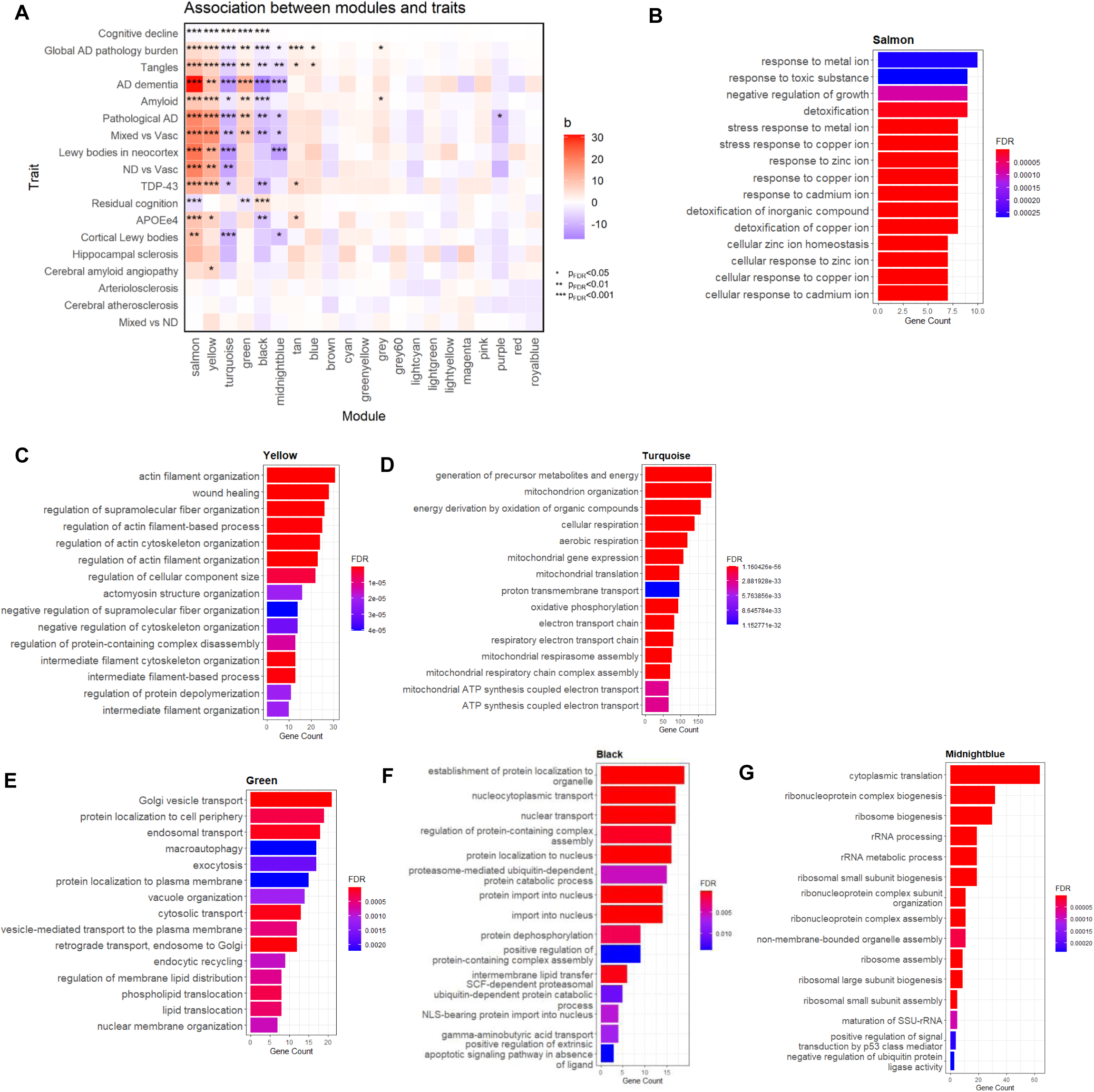
Proteomic co-expression network analysis identifies shared and distinct modules. (**A**) Module–trait relationships between module eigengenes and neuropathology groups and clinicopathological traits, with associations evaluated using regression models adjusted for age and sex. (**B–C**) Functional enrichment analyses of modules.

Two modules, salmon and yellow, exhibited increased eigengene expression in both neurodegenerative and mixed pathology relative to vascular pathology and showed strong positive associations with AD–related clinicopathological traits, including β-amyloid burden, tau tangle density, global AD pathology, TDP-43 pathology, neocortical Lewy bodies, pathological and clinical diagnoses of AD, and *APOE ε4* status, together with negative associations with cognitive decline and residual cognition (**Fig. 2A**). Functional enrichment analysis indicated that the salmon module was enriched for metal ion homeostasis and detoxification pathways, reflecting cellular responses to metabolic and toxic stress (**Fig. 2B**), whereas the yellow module was enriched for tissue remodeling and cytoskeletal reorganization processes, including wound healing and glial–vascular responses (**Fig. 2C**). To identify the proteins driving this shared enrichment, we examined the subset of pathway-contributing proteins common to both contrasts. These modules contained proteins implicated in glial activation, injury response, and AD risk, including reactive astrocyte markers (*GFAP, SDC4, CD44*), AD-linked proteins (*APP, CLU*), and stress- and tau-associated signaling mediators (*MAPK1/3, SMAD4*) (**Table S8, S9,** and **S10**). The co-occurrence of these proteins within modules upregulated in both neurodegenerative and mixed pathology is consistent with glial, immune, and cellular stress responses linked to AD pathology.

In contrast, the turquoise module showed reduced eigengene expression in both neurodegenerative and mixed pathology relative to vascular pathology and exhibited associations in the opposite direction to the salmon and yellow modules across clinicopathological traits, while associations with *APOE ε4* status and residual cognition were not significant (**Fig. 2A**). This module was strongly enriched for mitochondrial organization, oxidative phosphorylation, and respiratory chain pathways and was dominated by core bioenergetic and mitochondrial quality-control proteins, including mitochondrial import and cristae regulators (*TOMM40, IMMT, SAMM50*), respiratory chain components (*SDHA/SDHB, UQCRC1/2, COX6A1*), and metabolic gatekeepers (*PDHA1/PDHB*) (**Fig. 2D** and **Table S12**). Module–pathway concordance further supports coordinated regulation of immune and mitochondrial pathways rather than isolated protein-level effects.

Beyond the shared immune–metabolic core, additional co-expression modules showed selective associations with mixed pathology relative to vascular pathology, without corresponding changes in the neurodegenerative–vascular comparison. In particular, the green module exhibited increased eigengene expression in mixed pathology relative to vascular pathology (**Fig. 2A**) and was enriched for pathways related to vesicle-mediated transport, endosomal and Golgi trafficking, lipid and membrane dynamics, and macroautophagy (**Fig. 2E** and **Table S13**). Proteins within this module included adhesion and guidance molecules and regulators of vesicular trafficking, such as SLIT/ROBO family members and integrins, consistent with enhanced cellular transport and membrane remodeling programs in mixed pathology (**Table S10**). Conversely, the black and midnightblue modules showed reduced eigengene expression in mixed pathology relative to vascular pathology (**Fig. 2A**). Functional enrichment analysis indicated that the black module was enriched for nucleocytoplasmic transport, protein localization, ubiquitin–proteasome–mediated turnover, and proteostasis pathways (**Fig. 2F** and **Table S14**), whereas the midnightblue module was strongly enriched for ribosome biogenesis, rRNA processing, and cytoplasmic translation (**Fig. 2G** and **Table S15**). These patterns suggest selective attenuation of protein transport, quality control, and biosynthetic capacity in mixed pathology, distinguishing it from both vascular and neurodegenerative pathology alone.

### Regional transcriptomic architecture reveals a conserved immune–metabolic program

To characterize transcriptional programs associated with neurodegenerative, vascular, and mixed pathology across brain regions, we analyzed bulk RNA-seq data from the DLPFC, posterior cingulate cortex (PCC), and anterior caudate (AC). Differential gene expression analyses were conducted for three pairwise contrasts—neurodegenerative versus vascular, mixed versus vascular, and mixed versus neurodegenerative—adjusting for age, sex, and technical covariates. Across regions, single-transcript effects were modest: the mixed–vascular contrast yielded the greatest number of nominal DEGs, whereas no genes reached FDR significance in DLPFC for the mixed–neurodegenerative comparison, underscoring that bulk transcriptomic differences between pathology groups are distributed across coordinated gene sets rather than dominated by large effects at individual loci (**Fig. 3A-3C** and **Table S16-S18**).

**Fig. 3.**
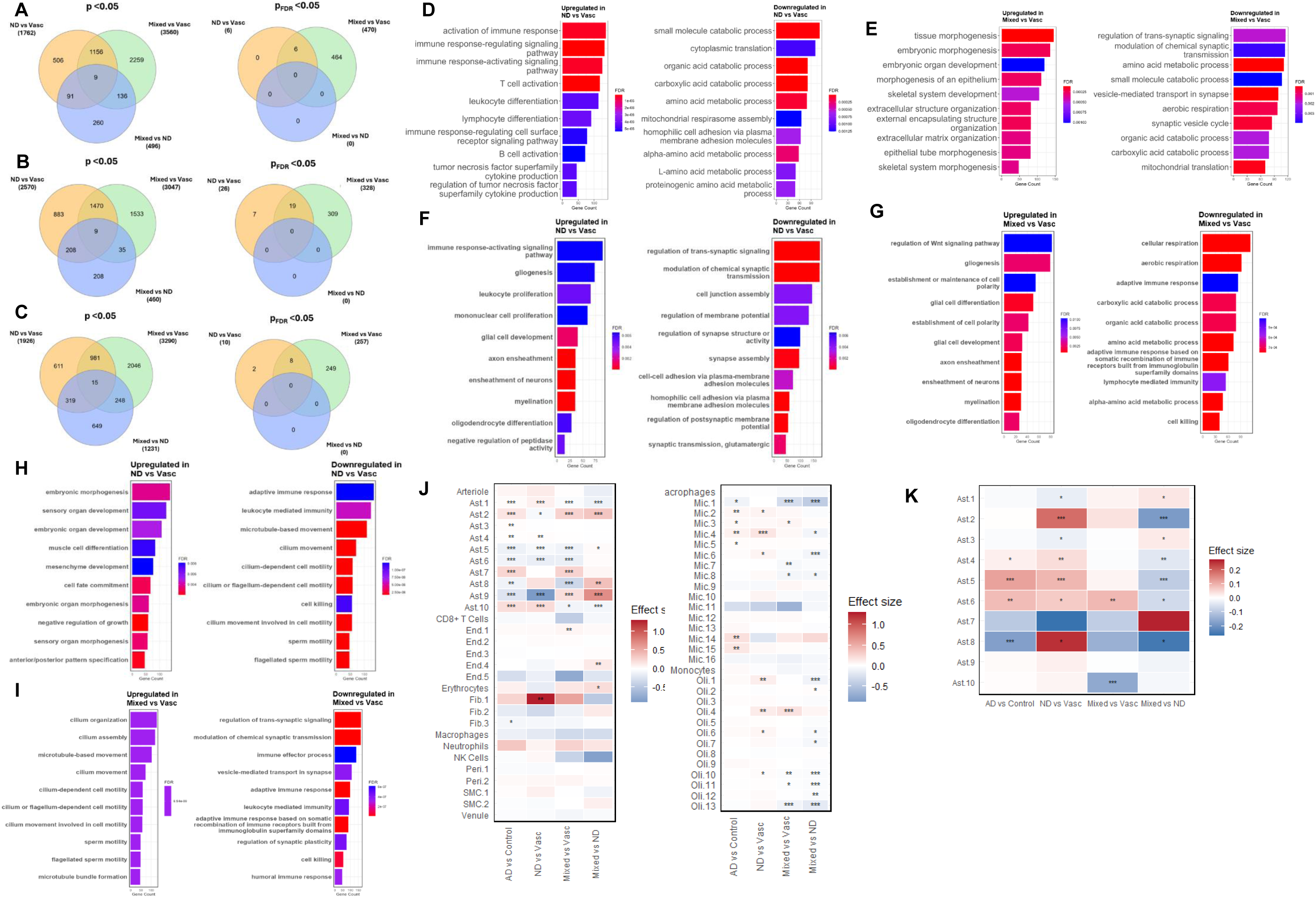
Regional transcriptomic analyses across brain regions. (**A–C**) Differential gene expression for neurodegenerative versus vascular, mixed versus vascular, and mixed versus neurodegenerative comparisons in DLPFC, PCC, and AC. (**D–I**) Gene Ontology enrichment analyses in DLPFC (D–E), PCC (F–G), and AC (H–I). (**J–K**) Cell-type–specific differential expression from single-nucleus RNA sequencing.

Despite modest gene-level signals, pathway analyses revealed a coherent transcriptional architecture. In DLPFC, neurodegenerative pathology relative to vascular pathology was characterized by upregulation of immune and inflammatory programs (e.g., immune activation and cytokine signaling) and downregulation of mitochondrial and bioenergetic pathways (e.g., oxidative phosphorylation and respiratory chain function) (**Fig. 3D** and **Table S19**). The mixed–vascular contrast recapitulated this immune–metabolic pattern, with additional enrichment of extracellular matrix and tissue remodeling processes (**Fig. 3E** and **Table S20**). Concordance between neurodegenerative–vascular and mixed–vascular contrasts was significant at the pathway level, particularly among downregulated pathways (Jaccard = 0.40, p = 0.038; **Table S6**).

This immune–metabolic transcriptional program extended beyond the DLPFC to additional brain regions. In both the neurodegenerative versus vascular (**Fig. 3F**, **Table S21** and **S22**) and mixed versus vascular contrasts (**Fig. 3G**, **Table S23** and **S24**), the PCC robustly recapitulated immune activation together with suppression of synaptic and metabolic pathways, whereas the AC exhibited weaker but directionally consistent effects (**Fig. 3H** and **3I**, **Table S25-S38**). Quantitative comparison of pathway enrichment across regions demonstrated significant regional concordance, with the strongest overlap observed between the DLPFC and PCC. This included concordant upregulation of immune pathways in the neurodegenerative versus vascular contrast (Jaccard = 0.21; p = 8.4 × 10⁻⁶), followed by intermediate concordance of downregulated pathways in the mixed versus vascular contrast (Jaccard = 0.38; p = 0.052), consistent with partial conservation of metabolic suppression (**Table S29**). The highest regional concordance was observed for pathways downregulated in the mixed versus neurodegenerative contrast (Jaccard = 0.55; p = 1.8 × 10⁻¹¹), which were dominated by immune effector and leukocyte-mediated processes (**Table S29**). In contrast, overlap involving the AC was limited across regions (**Table S29**), consistent with regional modulation of a shared transcriptional response.

Because bulk RNA-seq averages transcriptional signals across heterogeneous cellular populations, we next leveraged single-nucleus RNA-seq to resolve the cellular architecture underlying the attenuated transcript-level effects observed in bulk tissue. GFAP, which was robustly increased at the protein level in both neurodegenerative–vascular and mixed–vascular contrasts, exhibited consistent upregulation across multiple astrocyte subpopulations in single-nucleus data (**Fig. 3J**). This induction was distributed across several astrocytic states rather than confined to a single reactive cluster, indicating coordinated engagement of astrocyte programs. In contrast, APP demonstrated marked astrocyte-state–specific regulation, with variable direction and magnitude across astrocyte subpopulations (**Fig. 3K**).

These findings indicate that the conserved immune–metabolic signature observed across regions reflect coordinated but distributed transcriptional regulation, whose composite signal emerges at the pathway level in bulk tissue.

### Cross-omics convergence reveals conserved immune–metabolic programs

Having identified a shared immune activation and mitochondrial suppression signature within proteomic and transcriptomic datasets independently, we next evaluated whether these programs converge across molecular layers. We compared significantly enriched GO pathways (FDR < 0.05) between DLPFC proteomics and DLPFC RNA-seq for each pathology contrast and extended this analysis to transcriptomic data from the PCC and AC to assess regional robustness.

In the DLPFC, the neurodegenerative–vascular contrast demonstrated robust convergence across proteomic and transcriptomic layers (76 shared upregulated pathways; Jaccard = 0.23), predominantly involving leukocyte migration, chemotaxis, immune effector responses, angiogenesis, and wound-response programs (**Fig. 4A** and **Table S29**). Concordant downregulation converged on mitochondrial respiration and oxidative phosphorylation (8 shared pathways; Jaccard = 0.12), indicating coordinated suppression of bioenergetic capacity across molecular layers (**Fig. 4B** and **Table S29**). Thus, proteomic and transcriptomic data independently captured the same inflammatory activation coupled with metabolic attenuation.

**Fig. 4.**
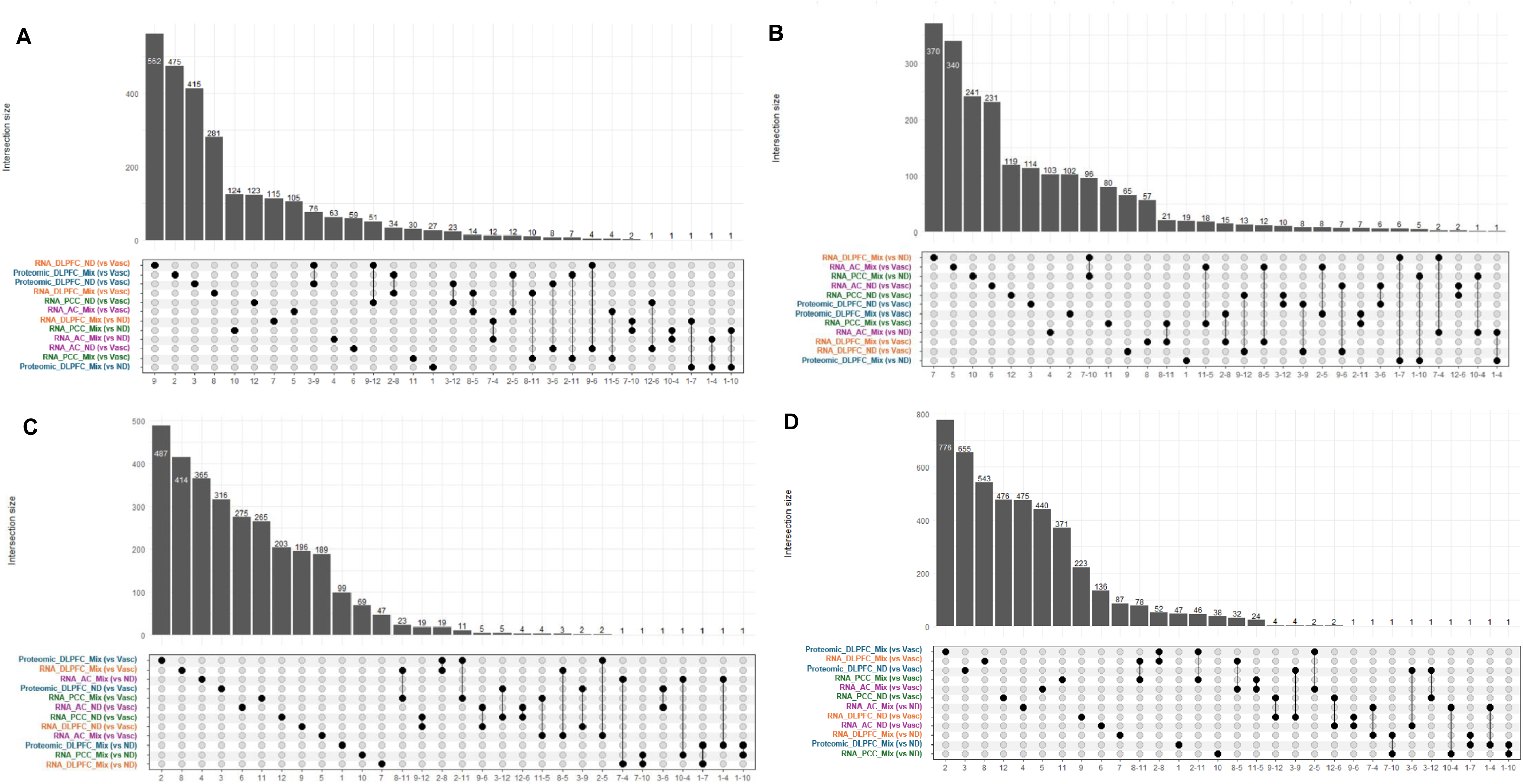
Cross-omics overlap of pathways and genes. (**A–B**) UpSet plots showing overlap of significantly enriched pathways across proteomic and transcriptomic datasets for each pathology comparison. (**C–D**) UpSet plots showing overlap of differentially expressed proteins and genes across datasets.

A similar immune–metabolic architecture was observed in the mixed–vascular contrast. Shared upregulated pathways (n = 34; Jaccard = 0.13) again reflected inflammatory and immune migration programs, while shared downregulated pathways (n = 15; Jaccard = 0.23) centered on mitochondrial respiratory and translational processes (**Fig. 4A** and **4B**, and **Table S29**). Although mixed pathology engages additional remodeling signatures within individual omics layers, the immune–metabolic axis remained the dominant cross-modal point of convergence.

In contrast, cross-omics overlap in the mixed–neurodegenerative comparison was minimal, consistent with the limited proteomic separation between these groups (**Table S29**). When DLPFC proteomics were compared with RNA-seq from PCC and AC, cross-omics concordance was attenuated but directionally preserved (Jaccard ∼0.04 –0.12), supporting partial regional conservation of this systems-level program (**Fig. 4A** and **4B**, and **Table S29**).

Importantly, this reproducibility at the pathway level contrasted with limited concordance at the level of individual genes and proteins across omics platforms (**Fig. 4C and 4D**). Although substantial gene-level overlap was observed between pathology contrasts within the proteomics dataset (e.g., Jaccard = 0.45–0.59) (**Table S30**), cross-modal overlap between proteomic and transcriptomic datasets remained low (generally ≤ 0.12), even when restricting to nominally significant features (p < 0.05) (**Table S31**). These findings indicate that cross-platform concordance is organized primarily at the level of coordinated biological programs rather than one-to-one transcript–protein correspondence.

These findings demonstrate that immune activation coupled with mitochondrial suppression as a conserved, cross-modal organizing principle of neurodegenerative and mixed pathology, conserved across molecular layers and broadly preserved across cortical regions.

## DISCUSSION

We provide a neuropathology-stratified molecular analysis of vascular, neurodegenerative, and mixed pathology in the aging human brain using matched proteomic and transcriptomic data from ROS/MAP. Across modalities and cortical regions, neurodegenerative and mixed pathology share a reproducible molecular signature defined by enrichment of immune and inflammatory pathways together with suppression of mitochondrial and oxidative phosphorylation programs relative to vascular pathology. This architecture was robust at the pathway level but showed limited correspondence at the level of individual genes and proteins.

The marked asymmetry between feature-level and program-level overlap is a central observation. Mixed and neurodegenerative groups showed minimal separation at the single-protein level, yet pathway analyses resolved coherent differences, including mixed-specific enrichment of extracellular matrix organization and vesicle-mediated transport processes. Similarly, cross-omics concordance was strongest for biological programs—immune migration and effector responses upregulated, mitochondrial respiration downregulated—whereas overlap among individually significant features remained sparse. These findings indicate that neuropathology-associated molecular organization is more consistently captured by coordinated pathway structure than by individual markers.

Proteomic network analysis reinforced this organization. Modules enriched for glial activation, stress responses, and tissue remodeling were elevated in neurodegenerative and mixed pathology and tracked established clinicopathologic indices, including amyloid and tau burden. In contrast, a mitochondrial module enriched for respiratory chain and bioenergetic proteins showed reduced expression in neurodegenerative and mixed groups but relative preservation in vascular pathology. These results define an immune–bioenergetic axis that distinguishes neurodegenerative-linked states from vascular-dominant pathology within the same community-based cohort.

These molecular findings align with clinicopathologic observations in the same cohorts. Neuropathologic stratification has operationally separated vascular, neurodegenerative, and mixed groups, with macroinfarcts—particularly in frontal white matter—associating with cognitive impairment in individuals without substantial neurodegenerative pathology, and high burdens of AD neuropathologic change and LATE-NC associating with the steepest longitudinal decline. Our data provide molecular context for these distinctions, demonstrating that vascular-dominant pathology is not simply a milder form of neurodegenerative remodeling and that mixed pathology is accompanied by additional structured remodeling beyond the shared inflammatory–mitochondrial program.

The pathway-level stability observed across omics layers has implications for biomarker interpretation. In the context of frequent comorbidity, cross-modal reproducibility was strongest for immune and mitochondrial programs rather than for individual analytes. This suggests that systems-level pathway signatures may offer more robust stratification across heterogeneous pathology than single molecular features.

Several limitations merit consideration. Neuropathology-defined groups impose categorical boundaries on continuous and overlapping processes. Bulk transcriptomic analyses cannot resolve cell-type–specific contributions to pathway signals, and regional variation indicates that conserved programs are modulated by anatomical context. Finally, the cross-sectional, postmortem design precludes inference regarding temporal sequence.

In summary, neuropathology-stratified multi-omics identifies a conserved immune activation and mitochondrial suppression program shared by neurodegenerative and mixed pathology relative to vascular pathology, together with additional remodeling signatures specific to mixed states. These results define a pathway-centered molecular framework for interpreting vascular and neurodegenerative contributions to late-life cognitive impairment.

## MATERIALS AND METHODS

### Study Design

Participants were enrolled from the Religious Orders Study (ROS) or the Rush Memory and Aging Project (MAP), two harmonized, prospective, community-based longitudinal cohort studies of aging and dementia. Both cohorts enroll participants aged 65 years or older who are free of known dementia at baseline and who agree to annual clinical and neuropsychological evaluations as well as postmortem organ donation (*14*). Detailed descriptions of the study design and cohort characteristics have been reported previously (*14*). Written informed consent and an Anatomic Gift Act were obtained from all participants. The study was approved by the Institutional Review Board of Rush University Medical Center.

To delineate vascular and neurodegenerative pathological signatures, participants were categorized into three neuropathology-defined subgroups—neurodegenerative (ND), vascular, and mixed—based on comprehensive postmortem pathological assessments, as previously described(*6*). A vascular subgroup was defined as participants without significant neurodegenerative brain pathologies, including intermediate or high likelihood of AD pathological diagnosis, extension of TDP-43 proteinopathy to the hippocampus or neocortex, hippocampal sclerosis, or neocortical Lewy body pathology. A neurodegenerative subgroup comprised participants without significant cerebrovascular disease pathologies, including macroinfarcts, microinfarcts, or moderate-to-severe atherosclerosis or arteriolosclerosis. Participants with moderate-to-severe cerebral amyloid angiopathy (CAA) were not excluded, given its close association with AD(*15*). The mixed subgroup included all remaining participants exhibiting overlapping neurodegenerative and vascular pathologies. To ensure clear separation of pathology-defined subgroups, participants whose pathological profiles did not meet subgroup criteria were excluded during data preprocessing. Specifically, individuals classified as neurodegenerative pathology without pathological AD (n=74) and those classified as vascular pathology with evidence of pathological AD (n=1) were removed prior to downstream analyses.

### RNA sequencing and data processing

RNA sequencing was performed across multiple batches, sequencing centers, and experimental protocols from postmortem human brain tissue from participants in the ROSMAP study, as previously described in detail (*16, 17*). For dorsolateral prefrontal cortex (DLPFC), RNA sequencing was initially conducted at the Broad Institute in 10 batches (n = 739). Total RNA was extracted using the Qiagen miRNeasy Mini Kit with the RNase-Free DNase Set, quantified by NanoDrop, and assessed for quality using the Agilent BioAnalyzer. Samples with RNA integrity number (RIN) > 5 and RNA yield ≥5 µg were used for library construction. Libraries were prepared using either a dUTP-based stranded protocol (8 batches) or a modified Illumina TruSeq protocol (2 batches) and sequenced on the Illumina HiSeq 2000 platform with 101 bp paired-end reads, targeting ∼50 million paired reads per sample. Additional DLPFC samples (n = 124, 2 batches) were sequenced at the New York Genome Center (NYGC) using the KAPA Stranded RNA-seq Kit with RiboErase and the Illumina NovaSeq 6000 platform (2 × 100 bp, ∼30 million reads per sample). Further samples (n = 229, single batch) were processed at the Rush Alzheimer’s Disease Center using the Chemagic RNA Tissue Kit, with RNA quality assessed by Fragment Analyzer, followed by TruSeq stranded library preparation and NovaSeq 6000 sequencing (2 × 150 bp, 40–50 million reads). RNA sequencing of anterior caudate (AC) and posterior cingulate cortex (PCC) was conducted across multiple sequencing centers using protocols matched to those applied for DLPFC. AC samples were sequenced at the Broad Institute and NYGC, while PCC samples were sequenced at the Broad Institute, NYGC, and the Rush Alzheimer’s Disease Center, using matched extraction, library preparation, and sequencing procedures.

Each brain region was preprocessed separately. Gene-level expression values were derived from RNA-seq data, and gene-level read count matrices were used to quantify expression. Counts were normalized using the trimmed mean of M-values (TMM) method and log2-transformed prior to modeling. Outlier samples were removed based on quantified expression profiles, and lowly expressed genes were filtered prior to analysis to minimize technical noise (*17*). After preprocessing and filtering, a total of 18,629 transcripts in DLPFC, 19,017 transcripts in PCC, and 17,574 transcripts in AC were retained for downstream analyses.

To account for demographic and technical sources of variation in gene expression, covariates for expression modeling were selected using a forward-selection strategy as previously described (*17*). For the DLPFC, selected covariates included age at death, sex, batch, library size, percentages of coding bases, aligned reads, ribosomal bases, UTR bases, intergenic bases, percentage duplication, median 5 prime to 3 prime bias, median 3 prime bias, median CV coverage, postmortem interval, and study index (ROS or MAP). For the PCC, selected covariates included age at death, sex, batch, library size, aligned reads, percentages of coding bases, ribosomal bases, UTR bases, percentage duplication, median 5 prime to 3 prime bias, median 3 prime bias, median CV coverage, postmortem interval, and study index. For the AC, selected covariates included age at death, sex, batch, aligned reads, ribosomal bases, usable bases, intergenic bases, percentage duplication, median 5 prime to 3 prime bias, median 5 prime bias, median CV coverage, postmortem interval, and study index. Gene expression values were adjusted for selected covariates using linear regression, with log2-transformed normalized expression as the outcome, and covariate-adjusted residuals were carried forward for downstream analyses. Residualized expression values represent transcript-level variation not explained by known demographic or technical factors.

Single-nucleus RNA sequencing data from DLPFC tissue of ROSMAP participants were obtained from a previously published dataset (*18*). Nuclei were isolated from frozen postmortem brain tissue and profiled to capture transcriptomic variation across brain cell populations. After quality control and filtering, data from 424 participants were retained for downstream analyses. Cell-type annotations provided by the original study (*18*), derived from unsupervised clustering and established marker genes, were used for all analyses. The annotated cell types included excitatory and inhibitory neurons, astrocytes, oligodendrocytes, microglia, and vascular and immune cell populations.

Association analyses were performed to compare pathological Alzheimer’s disease cases and controls, as well as three pairwise contrasts: neurodegenerative versus vascular, mixed versus vascular, and mixed versus neurodegenerative. For each gene and cell type, logistic regression models were fitted with pathology status as the dependent variable and gene expression as the predictor, adjusting for age at death and sex.

### Proteomic data processing

Proteomic profiling of DLPFC (Brodmann area 9) tissue from the ROSMAP cohort was performed using large-scale tandem mass tag (TMT)–based quantitative mass spectrometry, as described previously (*19*). Briefly, protein abundances were quantified following extensive sample randomization, TMT multiplexing, high-resolution LC–MS/MS, and database searching against the UniProt human proteome with stringent peptide- and protein-level false discovery rate control. Protein-level reporter ion intensities were assembled across TMT batches and subjected to rigorous normalization using the tunable median polish of ratios (TAMPOR) framework, which removes intra-batch, inter-batch, and inter-cohort technical variance while preserving biologically meaningful signal. This procedure includes transformation of reporter abundances to ratios followed by log2 scaling, normalization to appropriate within-batch reference medians, and iterative convergence across batches. Proteins with greater than 50% missing values were excluded, and no imputation was performed. As part of this processing, technical effects related to TMT batch structure were removed using the TAMPOR framework, and residual effects of postmortem interval (PMI) were subsequently adjusted by regression in the original study, yielding log2-scaled protein abundance estimates corrected for batch and PMI effects.

Using the fully processed DLPFC proteomic data generated by this pipeline, we analyzed all 10,030 quantified proteins. Protein abundance values were additionally adjusted for age at death and sex using linear regression models, with log2-scaled normalized protein abundance as the outcome variable, and the resulting covariate-adjusted residuals were used for all downstream analyses.

### Differential expression analysis

Differential expression analyses were performed separately for each omics modality using linear regression models applied to covariate-adjusted residuals. Three pairwise contrasts were evaluated for each dataset: neurodegenerative versus vascular pathology, mixed versus vascular pathology, and mixed versus neurodegenerative pathology. P-values were adjusted for multiple testing using the Benjamini–Hochberg procedure, with false discovery rate (FDR) < 0.05 considered statistically significant.

### Pathway enrichment analysis

To interpret differential molecular changes at the pathway level, Gene Set Enrichment Analysis (GSEA) (*20*) was performed using the gseGO function from the clusterProfiler R package (*21*) with Gene Ontology Biological Process (GO BP) gene sets. Pathways containing between 100 and 500 genes were tested for enrichment and pathways were considered significant at FDR < 0.05. For biological interpretation, we prioritized the top upregulated and downregulated GO BP pathways within each pairwise contrast, ranked by FDR within each direction.

To identify molecular drivers of enriched pathways, we examined genes that were both significantly differentially expressed (FDR < 0.05) and members of prioritized GO Biological Process pathways. To identify shared molecular contributors in the neurodegenerative versus vascular and mixed versus vascular contrasts, we examined genes belonging to the top 10 significantly upregulated and downregulated pathways in each comparison, ranked by FDR. Genes present in prioritized pathways across both contrasts were retained as a shared core set of pathway-contributing proteins. Differential protein expression statistics were reported for all three subgroup contrasts (ND vs Vasc, Mixed vs Vasc, and Mixed vs ND), and genes were annotated by weighted gene co-expression network module membership and predominant cell-type enrichment.

### Weighted gene co-expression network analysis (WGCNA)

To identify coordinated molecular programs beyond single-feature differential expression analyses, weighted gene co-expression network analysis (WGCNA) was performed on DLPFC proteomic data using the WGCNA R package (*22*). Signed co-expression networks were constructed based on pairwise Pearson correlations between gene expression profiles, and a soft-thresholding power was selected to approximate scale-free topology. Modules were identified using a minimum module size of 30 and a merge height of 0.4. Module eigengenes, defined as the first principal component of each module, were calculated and tested for association with pathology-defined subgroup contrasts. Module–trait association analyses were further conducted to assess relationships between module eigengenes and clinicopathological traits, including β - amyloid load, tau tangle density, global AD pathology burden, TDP-43 pathology, cortical and neocortical Lewy bodies, cerebral amyloid angiopathy, hippocampal sclerosis, cerebral atherosclerosis, arteriosclerosis, pathological diagnosis of AD, clinical diagnosis of AD dementia, cognitive decline estimated as the slope of global cognition derived from longitudinal assessments and residual cognition defined as the residuals of global cognition after adjustment for known brain pathologies. All associations were tested using generalized linear regression models adjusted for age and sex. Functional annotation of co-expression modules was performed using over-representation analysis of GO BP and Disease Ontology (DO) terms with clusterProfiler (*21*). Enrichment significance was assessed at FDR < 0.05.

### Cross-omics convergence analysis

To evaluate convergence of molecular signatures across omics layers, enrichment results were examined both within and across molecular modalities and brain regions. Pathway-level convergence was assessed as the primary measure of cross-omics consistency by comparing GO BP enrichment results derived from pairwise differential analyses of proteomic data from DLPFC and transcriptomic data from multiple brain regions including AC, DLPFC, and PCC. Pathways were considered convergent if they were significantly enriched (FDR < 0.05) in multiple omics modalities and exhibited concordant directionality, allowing assessment of whether subgroup-distinguishing biological programs were shared across regions or specific to individual regions. To further examine consistency at the gene level within convergent pathways, member genes contributing to each shared pathway were extracted and evaluated across omics layers. Analyses were restricted to genes that were significantly differentially expressed (FDR < 0.05) within each modality. Gene-level overlap analyses were conducted separately for genes derived from concordantly upregulated pathways and from concordantly downregulated pathways across omics modalities. Overlap between gene sets was quantified using Jaccard indices, and statistical significance was evaluated using Fisher’s exact tests.

Of note, all statistical analyses were performed using R 4.5.1 (https://www.r-project.org/).

## Supplementary Materials

Supplementary material is available with the online version of this article.

## Acknowledgements

We thank the participants of the Religious Orders Study and the Rush Memory and Aging Project, as well as the investigators and staff involved in data collection and curation.

## Funding

This work was supported by the National Institutes of Health (K01AG084849, U19AG078109, and P30AG066462 to A.J.L.), the National Alzheimer’s Coordinating Center–Alzheimer’s Association New Investigator Award (A.J.L.), and the Carol and Gene Ludwig Pilot Grant in Neurodegeneration (A.J.L.). ROSMAP was supported by National Institutes of Health (P30AG10161, R01AG15819, R01AG17917, and U01AG61356 to D.A.B.).

## Author contributions

A.J.L., R.M., and B.N.V. created the study concept and designed the study. A.J.L., M.L., E.Y., C.K., R.M., and B.N.V. interpreted the experiments. A.J.L., M.L., R.M., and B.N.V. drafted the manuscript. S.O., J.A.S., and D.A.B. provided ROSMAP data. All authors had access to the data and approved the final manuscript.

## Competing interests

The authors declare that they have no competing interests.

## Data and materials availability

The bulk RNA-seq data from three brain regions (dorsolateral prefrontal cortex, posterior cingulate cortex, and anterior caudate), as well as snRNA-seq and proteomics data generated by the Religious Orders Study and the Rush Memory and Aging Project, are available via the AD Knowledge Portal (https://adknowledgeportal.org). The AD Knowledge Portal is a platform for accessing data, analyses, and tools generated by the Accelerating Medicines Partnership for Alzheimer’s Disease (AMP-AD) Target Discovery Program and other National Institute on Aging (NIA)–supported programs to enable open-science practices and accelerate translational research. These data, analyses, and tools are shared early in the research cycle without a publication embargo on secondary use. Data are available for general research use in accordance with the Portal’s data access and attribution requirements (https://adknowledgeportal.org/DataAccess/Instructions). Processed RNA-seq data generated in this study are available on Synapse (Synapse: syn25741873).

## Notes

### Competing Interest Statement

The authors have declared no competing interest.

## References

1. M. D. Sweeney, A. Montagne, A. P. Sagare, D. A. Nation, L. S. Schneider, H. C. Chui, M. G. Harrington, J. Pa, M. Law, D. J. J. Wang, R. E. Jacobs, F. N. Doubal, J. Ramirez, S. E. Black, M. Nedergaard, H. Benveniste, M. Dichgans, C. Iadecola, S. Love, P. M. Bath, H. S. Markus, R. Al-Shahi Salman, S. M. Allan, T. J. Quinn, R. N. Kalaria, D. J. Werring, R. O. Carare, R. M. Touyz, S. C. R. Williams, M. A. Moskowitz, Z. S. Katusic, S. E. Lutz, O. Lazarov, R. D. Minshall, J. Rehman, T. P. Davis, C. L. Wellington, H. M. Gonzalez, C. Yuan, S. N. Lockhart, T. M. Hughes, C. L. H. Chen, P. Sachdev, J. T. O’Brien, I. Skoog, L. Pantoni, D. R. Gustafson, G. J. Biessels, A. Wallin, E. E. Smith, V. Mok, A. Wong, P. Passmore, F. Barkof, M. Muller, M. M. B. Breteler, G. C. Roman, E. Hamel, S. Seshadri, R. F. Gottesman, M. A. van Buchem, Z. Arvanitakis, J. A. Schneider, L. R. Drewes, V. Hachinski, C. E. Finch, A. W. Toga, J. M. Wardlaw, B. V. Zlokovic, Vascular dysfunction-The disregarded partner of Alzheimer’s disease. Alzheimers Dement 15, 158–167 (2019).

2. W. D. Brenowitz, C. D. Keene, S. E. Hawes, R. A. Hubbard, W. T. Longstreth, Jr., R. L. Woltjer, P. K. Crane, E. B. Larson, W. A. Kukull, Alzheimer’s disease neuropathologic change, Lewy body disease, and vascular brain injury in clinic- and community-based samples. Neurobiol Aging 53, 83–92 (2017).

3. A. Kapasi, C. DeCarli, J. A. Schneider, Impact of multiple pathologies on the threshold for clinically overt dementia. Acta Neuropathol 134, 171–186 (2017).

4. J. A. Schneider, Z. Arvanitakis, W. Bang, D. A. Bennett, Mixed brain pathologies account for most dementia cases in community-dwelling older persons. Neurology 69, 2197–2204 (2007).

5. L. Yu, T. Wang, L. Du, D. A. Bennett, J. A. Schneider, P. A. Boyle, Neuropathologic Profiles and Associated Cognitive Trajectories in Community-Living Older Adults. JAMA Netw Open 9, e2554354 (2026).

6. S. Oveisgharan, R. J. Dawe, L. Yu, A. Kapasi, K. Arfanakis, V. Hachinski, J. A. Schneider, D. A. Bennett, Frequency and Underlying Pathology of Pure Vascular Cognitive Impairment. JAMA Neurol 79, 1277–1286 (2022).

7. L. Y. Di Marco, A. Venneri, E. Farkas, P. C. Evans, A. Marzo, A. F. Frangi, Vascular dysfunction in the pathogenesis of Alzheimer’s disease - A review of endothelium-mediated mechanisms and ensuing vicious circles. Neurobiology of disease 82, 593–606 (2015).

8. K. Rannikmae, N. Samarasekera, N. A. Martinez-Gonzalez, R. Al-Shahi Salman, C. L. Sudlow, Genetics of cerebral amyloid angiopathy: systematic review and meta-analysis. Journal of neurology, neurosurgery, and psychiatry 84, 901–908 (2013).

9. J. C. Lambert, C. A. Ibrahim-Verbaas, D. Harold, A. C. Naj, R. Sims, C. Bellenguez, A. L. DeStafano, J. C. Bis, G. W. Beecham, B. Grenier-Boley, G. Russo, T. A. Thorton-Wells, N. Jones, A. V. Smith, V. Chouraki, C. Thomas, M. A. Ikram, D. Zelenika, B. N. Vardarajan, Y. Kamatani, C. F. Lin, A. Gerrish, H. Schmidt, B. Kunkle, M. L. Dunstan, A. Ruiz, M. T. Bihoreau, S. H. Choi, C. Reitz, F. Pasquier, C. Cruchaga, D. Craig, N. Amin, C. Berr, O. L. Lopez, P. L. De Jager, V. Deramecourt, J. A. Johnston, D. Evans, S. Lovestone, L. Letenneur, F. J. Moron, D. C. Rubinsztein, G. Eiriksdottir, K. Sleegers, A. M. Goate, N. Fievet, M. W. Huentelman, M. Gill, K. Brown, M. I. Kamboh, L. Keller, P. Barberger-Gateau, B. McGuiness, E. B. Larson, R. Green, A. J. Myers, C. Dufouil, S. Todd, D. Wallon, S. Love, E. Rogaeva, J. Gallacher, P. St George-Hyslop, J. Clarimon, A. Lleo, A. Bayer, D. W. Tsuang, L. Yu, M. Tsolaki, P. Bossu, G. Spalletta, P. Proitsi, J. Collinge, S. Sorbi, F. Sanchez-Garcia, N. C. Fox, J. Hardy, M. C. Deniz Naranjo, P. Bosco, R. Clarke, C. Brayne, D. Galimberti, M. Mancuso, F. Matthews, I. European Alzheimer’s Disease, Genetic, D. Environmental Risk in Alzheimer’s, C. Alzheimer’s Disease Genetic, H. Cohorts for, E. Aging Research in Genomic, S. Moebus, P. Mecocci, M. Del Zompo, W. Maier, H. Hampel, A. Pilotto, M. Bullido, F. Panza, P. Caffarra, B. Nacmias, J. R. Gilbert, M. Mayhaus, L. Lannefelt, H. Hakonarson, S. Pichler, M. M. Carrasquillo, M. Ingelsson, D. Beekly, V. Alvarez, F. Zou, O. Valladares, S. G. Younkin, E. Coto, K. L. Hamilton-Nelson, W. Gu, C. Razquin, P. Pastor, I. Mateo, M. J. Owen, K. M. Faber, P. V. Jonsson, O. Combarros, M. C. O’Donovan, L. B. Cantwell, H. Soininen, D. Blacker, S. Mead, T. H. Mosley, Jr., D. A. Bennett, T. B. Harris, L. Fratiglioni, C. Holmes, R. F. de Bruijn, P. Passmore, T. J. Montine, K. Bettens, J. I. Rotter, A. Brice, K. Morgan, T. M. Foroud, W. A. Kukull, D. Hannequin, J. F. Powell, M. A. Nalls, K. Ritchie, K. L. Lunetta, J. S. Kauwe, E. Boerwinkle, M. Riemenschneider, M. Boada, M. Hiltuenen, E. R. Martin, R. Schmidt, D. Rujescu, L. S. Wang, J. F. Dartigues, R. Mayeux, C. Tzourio, A. Hofman, M. M. Nothen, C. Graff, B. M. Psaty, L. Jones, J. L. Haines, P. A. Holmans, M. Lathrop, M. A. Pericak-Vance, L. J. Launer, L. A. Farrer, C. M. van Duijn, C. Van Broeckhoven, V. Moskvina, S. Seshadri, J. Williams, G. D. Schellenberg, P. Amouyel, Meta-analysis of 74,046 individuals identifies 11 new susceptibility loci for Alzheimer’s disease. Nat Genet 45, 1452–1458 (2013).

10. G. W. Beecham, K. Hamilton, A. C. Naj, E. R. Martin, M. Huentelman, A. J. Myers, J. J. Corneveaux, J. Hardy, J. P. Vonsattel, S. G. Younkin, D. A. Bennett, P. L. De Jager, E. B. Larson, P. K. Crane, M. I. Kamboh, J. K. Kofler, D. C. Mash, L. Duque, J. R. Gilbert, H. Gwirtsman, J. D. Buxbaum, P. Kramer, D. W. Dickson, L. A. Farrer, M. P. Frosch, B. Ghetti, J. L. Haines, B. T. Hyman, W. A. Kukull, R. P. Mayeux, M. A. Pericak-Vance, J. A. Schneider, J. Q. Trojanowski, E. M. Reiman, C. Alzheimer’s Disease Genetics, G. D. Schellenberg, T. J. Montine, Genome-wide association meta-analysis of neuropathologic features of Alzheimer’s disease and related dementias. PLoS Genet 10, e1004606 (2014).

11. H. K. Dong, J. A. Gim, S. H. Yeo, H. S. Kim, Integrated late onset Alzheimer’s disease (LOAD) susceptibility genes: Cholesterol metabolism and trafficking perspectives. Gene 597, 10–16 (2017).

12. J. M. Farfel, L. Yu, A. S. Buchman, J. A. Schneider, P. L. De Jager, D. A. Bennett, Relation of genomic variants for Alzheimer disease dementia to common neuropathologies. Neurology 87, 489–496 (2016).

13. Y. Zhao, C. X. Gong, From chronic cerebral hypoperfusion to Alzheimer-like brain pathology and neurodegeneration. Cellular and molecular neurobiology 35, 101–110 (2015).

14. D. A. Bennett, A. S. Buchman, P. A. Boyle, L. L. Barnes, R. S. Wilson, J. A. Schneider, Religious Orders Study and Rush Memory and Aging Project. J Alzheimers Dis 64, S161–S189 (2018).

15. P. A. Boyle, L. Yu, S. Nag, S. Leurgans, R. S. Wilson, D. A. Bennett, J. A. Schneider, Cerebral amyloid angiopathy and cognitive outcomes in community-based older persons. Neurology 85, 1930–1936 (2015).

16. A. J. Lee, Y. Ma, L. Yu, R. J. Dawe, C. McCabe, K. Arfanakis, R. Mayeux, D. A. Bennett, H. U. Klein, P. L. De Jager, Multi-region brain transcriptomes uncover two subtypes of aging individuals with differences in Alzheimer risk and the impact of APOEepsilon4. bioRxiv, (2023).

17. L. Yu, S. Tasaki, J. A. Schneider, K. Arfanakis, D. M. Duong, A. P. Wingo, T. S. Wingo, N. Kearns, G. R. J. Thatcher, N. T. Seyfried, A. I. Levey, P. L. De Jager, D. A. Bennett, Cortical Proteins Associated With Cognitive Resilience in Community-Dwelling Older Persons. JAMA Psychiatry 77, 1172–1180 (2020).

18. A. Cain, M. Taga, C. McCabe, G. S. Green, I. Hekselman, C. C. White, D. I. Lee, P. Gaur, O. Rozenblatt-Rosen, F. Zhang, E. Yeger-Lotem, D. A. Bennett, H. S. Yang, A. Regev, V. Menon, N. Habib, P. L. De Jager, Multicellular communities are perturbed in the aging human brain and Alzheimer’s disease. Nat Neurosci 26, 1267–1280 (2023).

19. E. C. B. Johnson, E. K. Carter, E. B. Dammer, D. M. Duong, E. S. Gerasimov, Y. Liu, J. Liu, R. Betarbet, L. Ping, L. Yin, G. E. Serrano, T. G. Beach, J. Peng, P. L. De Jager, V. Haroutunian, B. Zhang, C. Gaiteri, D. A. Bennett, M. Gearing, T. S. Wingo, A. P. Wingo, J. J. Lah, A. I. Levey, N. T. Seyfried, Large-scale deep multi-layer analysis of Alzheimer’s disease brain reveals strong proteomic disease-related changes not observed at the RNA level. Nat Neurosci 25, 213–225 (2022).

20. A. Subramanian, P. Tamayo, V. K. Mootha, S. Mukherjee, B. L. Ebert, M. A. Gillette, A. Paulovich, S. L. Pomeroy, T. R. Golub, E. S. Lander, J. P. Mesirov, Gene set enrichment analysis: a knowledge-based approach for interpreting genome-wide expression profiles. Proc Natl Acad Sci U S A 102, 15545–15550 (2005).

21. T. Wu, E. Hu, S. Xu, M. Chen, P. Guo, Z. Dai, T. Feng, L. Zhou, W. Tang, L. Zhan, X. Fu, S. Liu, X. Bo, G. Yu, clusterProfiler 4.0: A universal enrichment tool for interpreting omics data. Innovation (Camb) 2, 100141 (2021).

22. P. Langfelder, S. Horvath, WGCNA: an R package for weighted correlation network analysis. BMC Bioinformatics 9, 559 (2008).

