## Supplemental Figures for "Multi-omics characterization of vascular, neurodegenerative, and mixed neuropathology in the aging human brain"

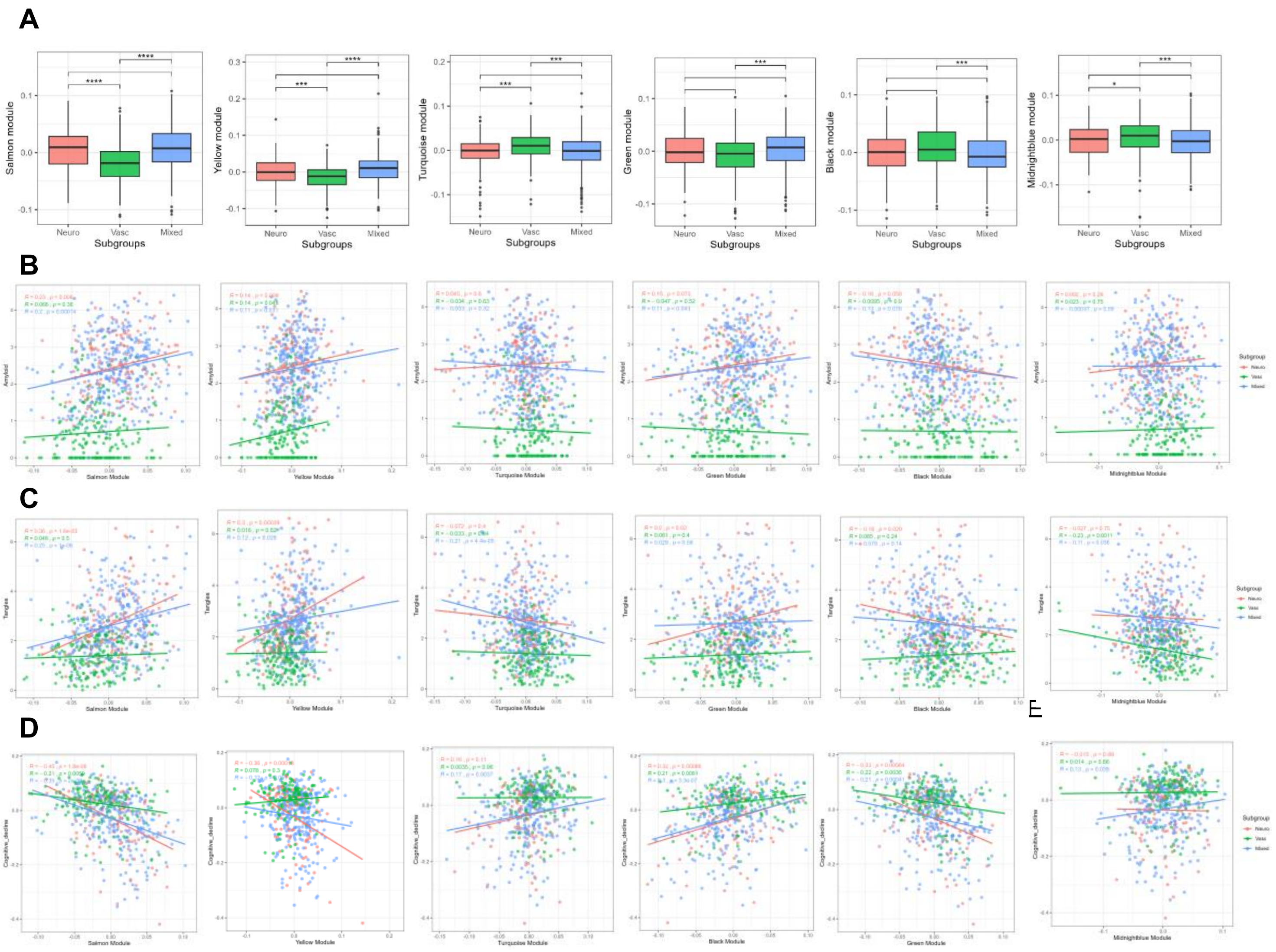


**Fig. S1. Proteomic co-expression modules and associations with clinicopathological traits.** (A) Comparison of module eigengene expression across neuropathology subgroups. (**B–D**) Associations of module eigengenes with β-amyloid burden (B), tau tangle density (C), and cognitive decline (D). Associations were evaluated using regression models adjusted for age and sex.
